# Functions and trafficking mechanisms of RIC-8 in *C. elegans* and mammalian cilia

**DOI:** 10.64898/2026.02.07.704597

**Authors:** Christina M. Campagna, Abigail E. Descoteaux, Abigail Poole, Eric Peet, Nawaphat Malaiwong, Michael P. O’Donnell, Inna Nechipurenko

**Affiliations:** Department of Biology and Biotechnology, Worcester Polytechnic Institute, Worcester, MA, USA; Department of Molecular, Cellular and Developmental Biology, Yale University, New Haven, CT, USA

## Abstract

Primary cilia exhibit conserved organization and contain structural and functional domains of unique molecular composition. The inversin compartment (InvC), which is found in the proximal ciliary segment of a subset of vertebrate and invertebrate cell types, concentrates different classes of signaling molecules. Mutations in genes encoding resident proteins of the InvC manifest in ciliopathies, highlighting the importance of the InvC in cilia biology. We previously showed that a chaperone of Gα proteins RIC-8 localizes to the InvC of *C. elegans* channel cilia; however, the mechanisms that regulate RIC-8 targeting to this ciliary sub-domain or RIC-8 function in the InvC remain unknown. Here, we build on our prior work to demonstrate that RIC-8 becomes restricted to the InvC during larval development and show that, while the RVxP motif and intact transition zone are required for its proper intraciliary distribution, RIC-8 localization to the cilium depends on intraflagellar transport. Using the ASH neuron as a model, we establish that RIC-8 modulates chemosensory responses mediated by channel cilia. Finally, we show that human RIC8A and RIC8B proteins are required for ciliogenesis in RPE-1 cells. Collectively, our results define ciliary trafficking mechanisms and novel functions for a highly conserved signaling protein.

## INTRODUCTION

Primary cilia are signaling organelles that extend from the surface of most mammalian cells and invertebrate sensory neurons (Anvarian et al., 2019; Derderian et al., 2023). The signaling capacity of primary cilia depends on compartmentalized localization of signaling proteins to distinct sub-ciliary domains. For example, the inversin compartment (InvC), defined by localization of INVS/NPHP-2, is found in the proximal primary cilium of many cell types (e.g. (Bennett et al., 2020; Nakajima et al., 2018; Shiba et al., 2009; Warburton-Pitt et al., 2014; Watanabe et al., 2003)) and concentrates cyclic nucleotide-gated channels (Nechipurenko and Sengupta, 2025; Wojtyniak et al., 2013), the small GTPase ARL-13 (Cevik et al., 2013), and the Gα protein chaperone RIC-8 (Campagna et al., 2023). Likewise, the TRPV channel and CDKL5 kinase are sequestered in the proximal regions of mechanosensory cilia in *Drosophila* (Xiang et al., 2022) and *Chlamydomonas* flagella (Tam et al., 2013), respectively. In addition to the InvC, the cilia tip concentrates select signaling proteins such as Hedgehog pathway components in mammalian cells (Haycraft et al., 2005; Wen et al., 2010) and receptor guanylate cyclase GCY-22 in the *C. elegans* ASER neuron (van der Burght et al., 2020).

Although the mechanisms that target specific signaling proteins to discrete ciliary sub-domains are not well understood, intraflagellar transport (IFT) – a conserved active transport system that carries many transmembrane and soluble proteins in and out of the cilium – has been reported to participate in localizing Gli proteins (Haycraft et al., 2005; Qin et al., 2011), GCY-22 (van der Burght et al., 2020), and ARL-13 (Cevik et al., 2013) to their respective sub-ciliary domains. IFT is organized into IFT-A and IFT-B multi-protein complexes that function as adapters between motors and ciliary protein cargoes. Kinesin-2 and dynein-2 motors drive IFT along ciliary microtubules in the anterograde (from cilia base to tip) and retrograde (from cilia tip to base) directions, respectively. In *C. elegans*, two kinesin-2 family motors (kinesin-II and OSM-3) work together to build cilia and mediate anterograde transport (Snow et al., 2004).

In *C. elegans*, cilia extend from the distal dendrites of a subset of sensory neurons and display distinct morphologies and molecular composition (Inglis et al., 2007; Nechipurenko and Sengupta, 2025). Most chemosensory neurons of the amphid and phasmid sensory organs in the worm head and tail, respectively, possess rod-like ‘channel’ cilia, while three types of amphid olfactory neurons (AWA, AWB, and AWC) have elaborate ‘wing’ cilia (Doroquez et al., 2014; Perkins et al., 1986; Ward et al., 1975). Although core ciliogenic mechanisms are conserved, cell-specific functions for cilia proteins are being increasingly observed in worms and mammals (e.g. (Campagna et al., 2023; Ditirro et al., 2019; Lewis et al., 2019; Rachel et al., 2012; Roayaie et al., 1998; Salama et al., 2025).

We previously reported that RIC-8, a cytoplasmic guanine nucleotide exchange factor and chaperone for heterotrimeric Gα protein subunits, shapes wing-cilia morphology by regulating levels of Gα ODR-3 (Campagna et al., 2023). Although RIC-8 is largely dispensable for the assembly of channel cilia, it localizes to the InvC of this cilia type (Campagna et al., 2023). Here, we demonstrate that RIC-8 modulates sensory responses mediated by ASH channel cilia, define molecular mechanisms that regulate its localization to this cilia type, and show that human RIC8A and RIC8B contribute to ciliogenesis in RPE-1 cells. Overall, our findings uncover new cilia roles for an evolutionarily conserved signaling protein.

## RESULTS AND DISCUSSION

### RIC-8 becomes restricted to the InvC during larval development independently of NPHP-2

When expressed from a multicopy transgene, RIC-8 localizes to the InvC of channel cilia in adult hermaphrodites (Campagna et al., 2023). We confirmed this localization pattern by tagging the endogenous *ric-8* locus with a split-Scarlet reporter (split-Sc) (Goudeau et al., 2021). Specifically, we inserted the 11^th^ β strand of codon-optimized Scarlet (*Scarlet_11_*) before the stop codon of *ric-8* and expressed the remaining *Scarlet_1-10_*fragment under the *bbs-8* promoter to reconstitute fluorescence and thus visualize localization of RIC-8::split-Sc in all ciliated neurons. Consistent with our earlier findings, RIC-8::split-Sc was distributed throughout phasmid neurons and exhibited enrichment in proximal cilia segments (Figure 1A) (Campagna et al., 2023).

**Figure 1.**
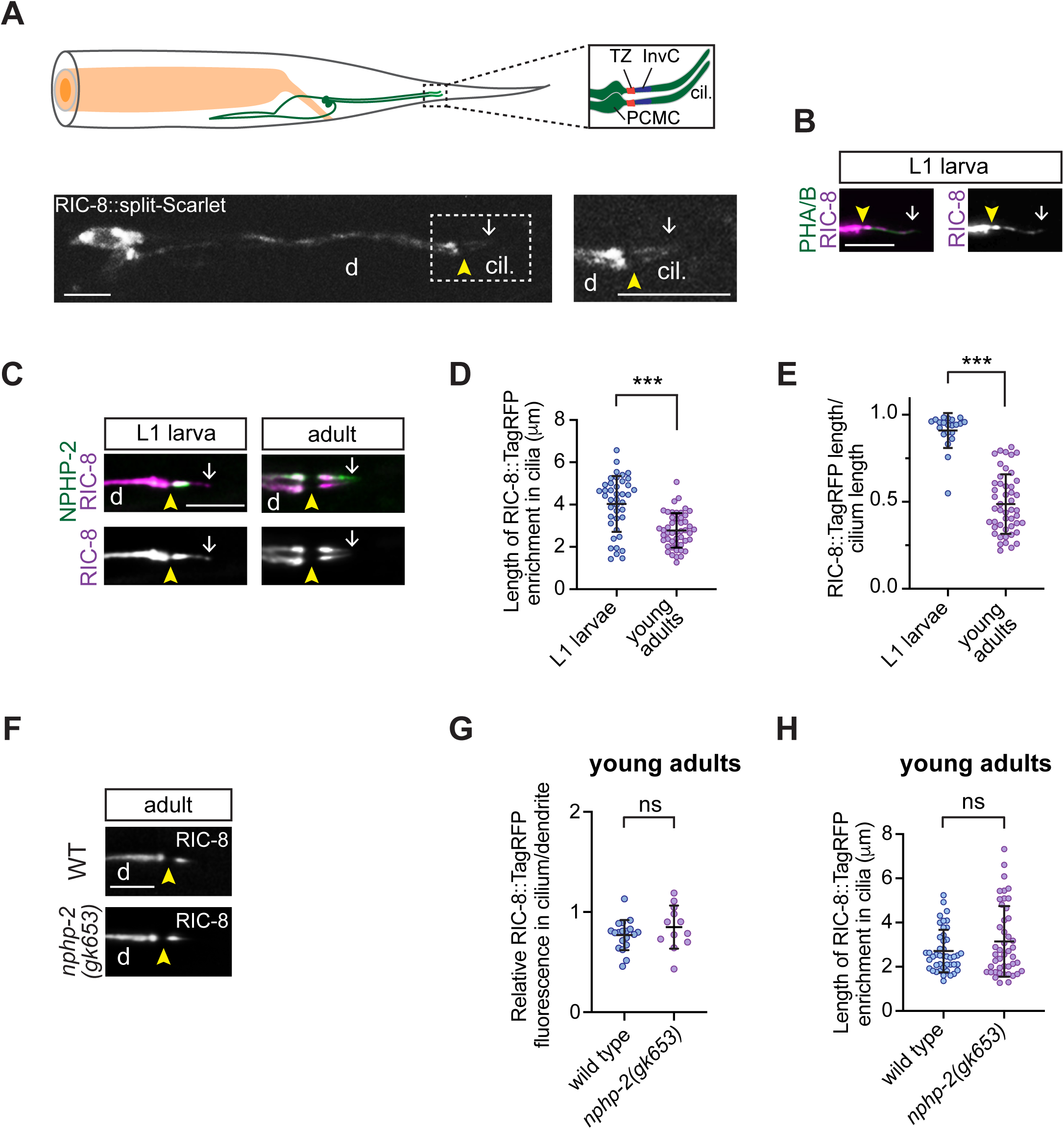
RIC-8 becomes restricted to the InvC during larval development. (A) *Top*: diagram of phasmid neurons in the worm tail. Phasmid cilia are magnified in the inset on the right. PCMC: periciliary membrane compartment; TZ: transition zone; InvC: inversin compartment. *Bottom*: RIC-8::split-Sc in phasmids. Boxed region is shown at higher magnification on the right. d: dendrite; cil.: cilia. (B and C) Localization of RIC-8::TagRFP (B) and co-localization of RIC-8::TagRFP with NPHP-2::GFP (C) in WT phasmids at the indicated stages. (D and E) Quantification of the absolute RIC-8::TagRFP signal length (D) and RIC-8::TagRFP signal length relative to phasmid cilia length (E) in L1 and adult WT animals. *** Different from L1 at p<0.001 (Mann-Whitney test). (F – H) Images (F) and quantification of relative RIC-8::TagRFP fluorescence (G) and RIC-8::TagRFP signal length (H) in phasmid cilia of WT and *nphp-2* mutants. ns: not significant (Welch’s t test) (G), (Mann-Whitney test) (H). Arrowheads: TZ; arrows: distal boundary of RIC-8::TagRFP signal. Scale: 5 μm.

The InvC in phasmid cilia is established before the first-larval (L1) stage (Warburton-Pitt et al., 2014); however, at least one InvC resident protein (ARL-13) does not become restricted to the proximal cilium until L3 larval stage (Cevik et al., 2013). In contrast, *Drosophila* TRPV subunit Iav localizes directly to the proximal zone during cilia assembly and remains restricted there during extension of distal cilia segments (Xiang et al., 2022). To determine the timing of RIC-8 localization to the InvC, we analyzed a published transgenic strain that expresses RIC-8::TagRFP in all ciliated neurons (Campagna et al., 2023) due to rapid bleaching of RIC-8::split-Sc. Unlike in adults, RIC-8::TagRFP signal in L1 larvae extended outside the InvC, visualized with NPHP-2::GFP, into distal cilia segments (Figure 1, B – E), indicating that RIC-8 becomes restricted to the InvC during larval development similarly to ARL-13. Relative ciliary levels and intra-ciliary distribution of RIC-8::TagRFP were comparable in wild type (WT) and *nphp-2* mutants (Figure 1, F – H), suggesting that NPHP-2 does not regulate RIC-8 localization to or within cilia.

### The transition zone regulates intraciliary distribution of RIC-8 in channel cilia

The transition zone (TZ) functions as a selective barrier at the cilia base, and defects in TZ integrity are associated with altered localization of ciliary proteins including InvC components (Barbelanne et al., 2024; Cevik et al., 2013; Jensen et al., 2015; Li et al., 2016). To determine whether the TZ contributes to RIC-8 ciliary import or its restriction to the InvC, we examined localization of RIC-8::TagRFP in phasmid cilia of *mks-5(tm3100)* mutants with severely compromised TZ (Li et al., 2016; Schouteden et al., 2015). RIC-8::TagRFP levels inside the cilium relative to the distal dendrite were significantly reduced in *mks-5* mutants compared to WT (Figure 2, A and B), suggesting the TZ may contribute to RIC-8 retention inside the cilium.

**Figure 2.**
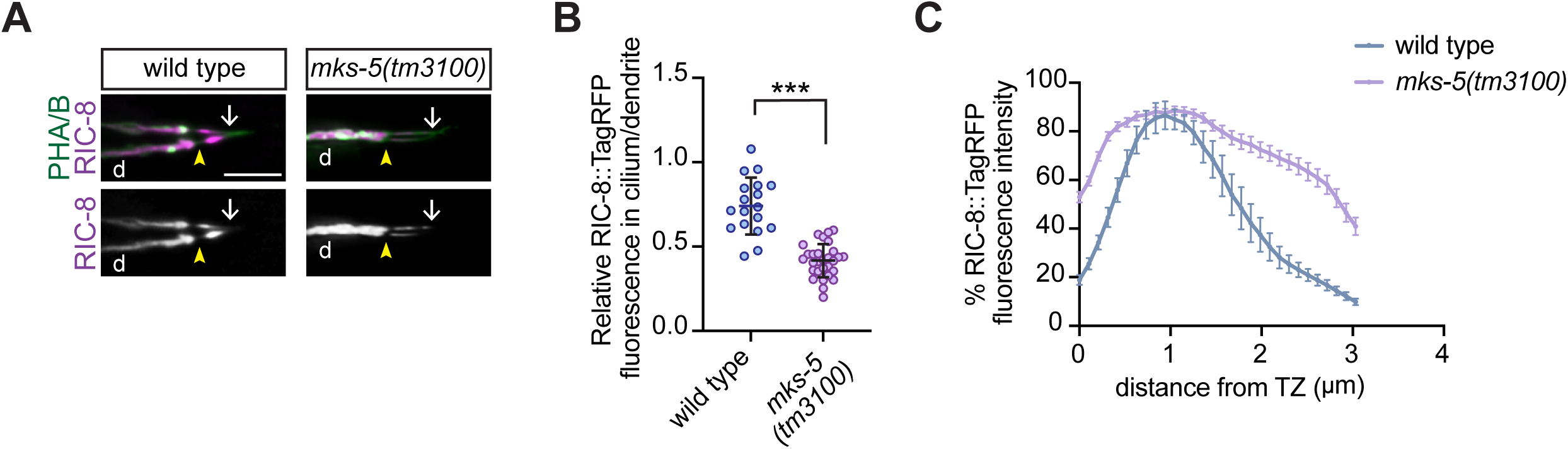
RIC-8 cilia localization depends on an intact transition zone. (A and B) Images (A) and quantification (B) of RIC-8::TagRFP localization in WT and *mks-5(tm3100)* adult phasmids. Arrowheads: TZ; arrows: distal boundary of RIC-8::TagRFP signal; d: dendrite. Scale: 5 μm. *** Different from wild type at p<0.001 (Welch’s t test). (C) Line scans of RIC-8::TagRFP intensity in phasmid cilia of the indicated genotypes. Zero corresponds to the cilium base. Error bars: SEM. n=19 and 36 neurons for WT and *mks-5(tm3100)*, respectively.

Indeed, other studies reported ectopic accumulation of cilia signaling proteins in the distal dendrite of TZ mutants, proposing that the TZ prevents proteins from ‘leaking’ out of the cilium (Brear et al., 2014; Cevik et al., 2013; Jensen et al., 2015; Li et al., 2016).

In addition to changes in the relative levels, we noted altered distribution of RIC-8::TagRFP inside *mks-5* mutant cilia. In WT, RIC-8::TagRFP signal peaks around 1 μm distal to the TZ (Figure 2, A and C) (Campagna et al., 2023), while in *mks-5* mutants it is more uniformly distributed throughout the proximal cilium (Figure 2, A and C). Our results suggest that TZ integrity is important for RIC-8 cilia retention and proper sub-ciliary distribution. The defects in RIC-8::TagRFP distribution in *mks-5* mutant are likely due to mislocalization of cilia proteins that restrict RIC-8 to the InvC.

### The RVxP motif helps restrict RIC-8 to the InvC

The RVxP motif plays a role in targeting select proteins to cilia (Deretic et al., 2005; Geng et al., 2006; Higginbotham et al., 2012; Jenkins et al., 2006; Mariani et al., 2016; Zuo et al., 2019). For example, mutations in the RVxP motif of the mammalian Arl13b exclude mutant protein from cilia (Higginbotham et al., 2012), while *C. elegans* ARL-13 with deleted RVxP exhibits expanded localization to distal cilia (Cevik et al., 2013). RIC-8 contains the RVIP motif in the carboxyl terminus (Supplemental Figure S1A), so we wanted to test whether this motif is necessary for either targeting RIC-8 to the cilium or restricting it to the InvC. Unlike WT RIC-8, RIC-8^ΔRVIP^ extended into distal ciliary segments (Supplemental Figure S1, B and C), pointing to a role for RVIP in sequestering RIC-8 in the InvC, similarly to the RVxP motif in ARL-13 (Cevik et al., 2013). It will be important to determine whether these changes in intraciliary RIC-8 distribution are accompanied by deficits in cilia function.

### RIC-8 ciliary localization is dependent on IFT

Next, we tested whether IFT plays a role in RIC-8 ciliary localization. In *C. elegans* channel cilia, the heterotrimeric kinesin-II motor moves anterograde IFT trains across the TZ (Mitra et al., 2024; Prevo et al., 2015; Snow et al., 2004). We examined RIC-8::TagRFP localization in *kap-1* and *klp-11* kinesin-II subunit mutants and noted significant reduction in RIC-8::TagRFP levels inside phasmid channel cilia relative to the periciliary membrane compartment (PCMC) in the distal dendrite (Figure 3, A and B), suggesting that the kinesin-II motor is required for localizing RIC-8 to the cilium. To determine whether the dynein-2 motor, which powers retrograde IFT trains, is important for trafficking RIC-8 out of the cilium, we investigated mutants of *che-3* and *xbx-1* that encode dynein heavy and light intermediate chains, respectively. Both mutants exhibited an increase in relative intraciliary RIC-8::TagRFP levels compared to WT (Figure 3, A and B). Together, these findings suggest that RIC-8 localization to the cilium depends on kinesin-II and dynein-2 motors.

**Figure 3.**
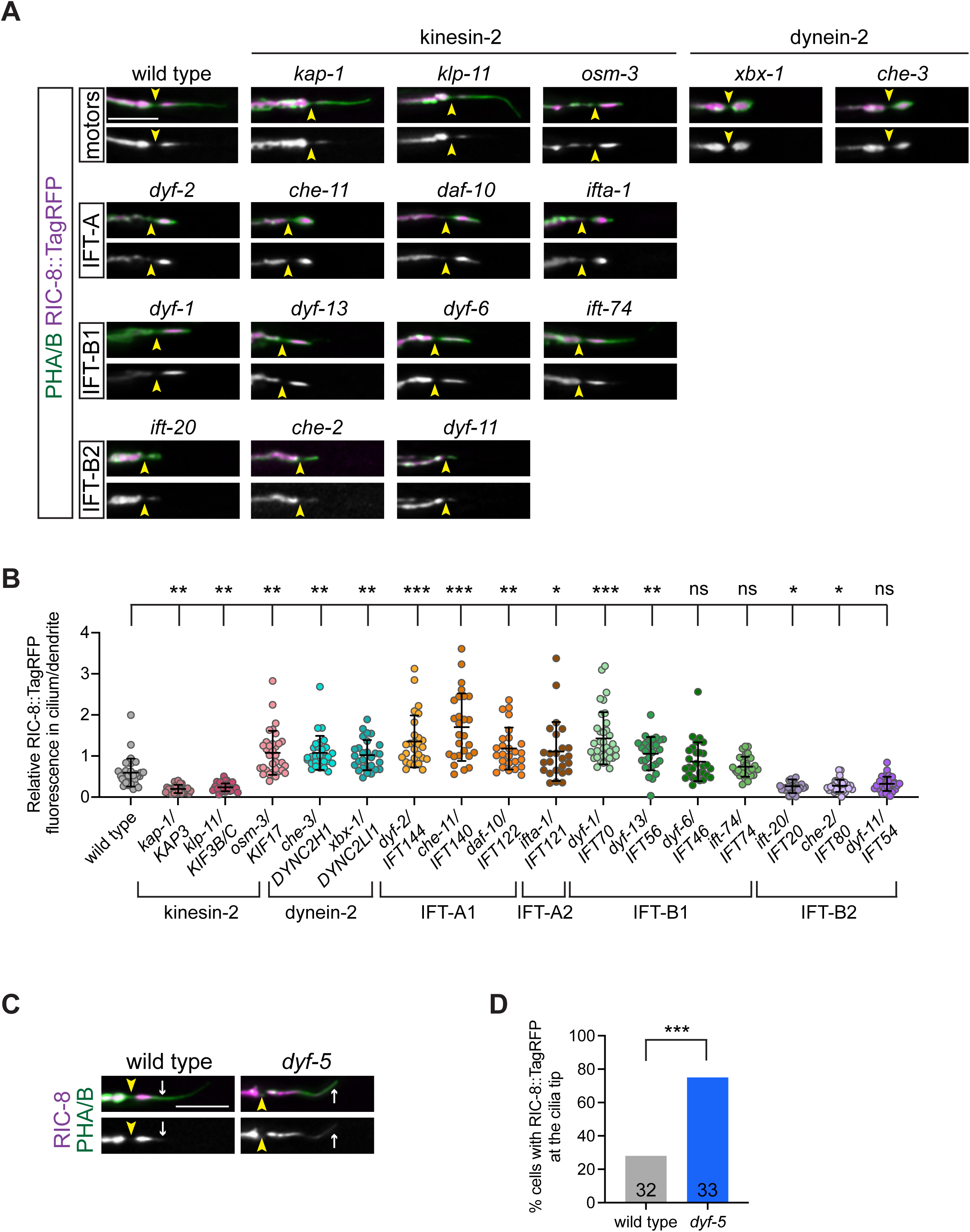
RIC-8 localization to channel cilia depends on IFT. (A and B) Images (A) and quantification (B) of RIC-8::TagRFP localization in phasmids of the indicated genotypes. Corresponding mammalian homologs are listed on the x-axis in (B). (C and D) Images (C) and quantification (D) of RIC-8::TagRFP localization in WT and *dyf-5(ok1177)* phasmids. Arrowheads: TZ; arrows: distal boundary of RIC-8::TagRFP signal. Scale: 5 μm. *, **, and *** Different from WT at p<0.05, 0.01, and 0.001, respectively (Kruskal-Wallis with Dunn’s multiple comparisons test) (B) and Fisher’s exact test (D).

The IFT-B complex comprised of the IFT-B1 and IFT-B2 sub-complexes mediates ciliary import and/or anterograde transport of several soluble proteins (Ahmed et al., 2008; Bhogaraju et al., 2013; Dai et al., 2018; Hou and Witman, 2017; Hunter et al., 2018; Ishikawa et al., 2014; Lacey and Pigino, 2024; Nakamura et al., 2020; Taschner et al., 2017; Zhao et al., 2020). On the other hand, IFT-A, which similarly consists of two sub-complexes (IFT-A1 and IFT-A2) plays a major role in retrograde transport (Blacque et al., 2006; Engel et al., 2012; Iomini et al., 2009; Piperno et al., 1998). To determine if RIC-8 trafficking in and out of the cilium depends on IFT-B and/or IFT-A, respectively, we quantified relative RIC-8::TagRFP levels in cilia of a subset of IFT-A/B gene mutants. Mutations in genes encoding IFT-A1 (*dyf-2*, *che-11*, and *daf-10*) and IFT-A2 (*ifta-1*) proteins increased accumulation of RIC-8::TagRFP inside the cilium relative to the PCMC, similarly to our observations in *che-3* and *xbx-1* dynein-2 mutants (Figure 3, A and B). These findings are consistent with the hypothesis that retrograde IFT contributes to RIC-8 transport out of the cilium. On the other hand, mutations in the IFT-B2 complex genes *ift-20* and *che-2* significantly reduced the cilium/PCMC ratio of RIC-8::TagRFP (Figure 3, A and B). *dyf-11* mutants showed a similar, although not statistically significant, decrease in relative RIC-8::TagRFP levels inside their cilia (Figure 3, A and B). These results suggest that ciliary localization of RIC-8 depends on at least a subset of IFT-B2 proteins. Furthermore, since all examined mutations in the IFT-B2, IFT-A, *osm-3*, and dynein-2 genes truncate cilia yet exert distinct effects on relative RIC-8 levels (Figure 3A), the observed changes in RIC-8 localization are unlikely to be simply a consequence of shorter cilia length.

Interestingly, mutations in the IFT-B1 genes *dyf-1* and *dyf-13* significantly increased ciliary RIC-8::TagRFP levels relative to the PCMC, while those in *dyf-6* and *ift-74* had no significant impact (Figure 3, A and B). Previous studies in *C. elegans* proposed that DYF-1 and possibly DYF-13 are required for activation of the homodimeric kinesin-2 motor OSM-3 and/or its loading onto the IFT-B module in channel cilia (Ou et al., 2005; Ou et al., 2007). This model was based on the observation that *dyf-1* mutants had no detectable OSM-3 transport inside the cilia; however, other IFT-A and IFT-B1/B2 components exhibited normal motility presumably due to being transported by heterotrimeric kinesin-II (Ou et al., 2005; Ou et al., 2007). Consistently, we find that *osm-3* mutants exhibited increased RIC-8::TagRFP fluorescence in cilia relative to the PCMC similarly to *dyf-1* and *dyf-13* mutant animals (Figure 3, A and B). These results suggest that OSM-3 and IFT-B1 components DYF-1 and DYF-13 may participate in ciliary trafficking of RIC-8.

A recent study in *Chlamydomonas* demonstrated that anterograde IFT trains undergo extensive remodeling at the cilia tip into retrograde trains of distinct conformation (Lacey and Pigino, 2024). The rearrangement of IFT-A/B components during this remodeling event generates unique cargo-binding interfaces in anterograde vs retrograde trains. Notably, IFT70/DYF-1 was proposed to form a potential cargo-binding surface on retrograde IFT trains. Thus, it would be interesting to determine whether RIC-8 is carried on retrograde trains via binding to DYF-1, DYF-13, or OSM-3, which is also moved by retrograde IFT from the cilia tip to the middle segment.

### DYF-5 restricts RIC-8 to the InvC

The MAK/ICK kinase DYF-5 regulates cilia length and IFT protein localization in *C. elegans* sensory neurons (Burghoorn et al., 2007; Maurya et al., 2019; Mul et al., 2025; Yi et al., 2018). *dyf-5* mutants exhibit long cilia and ectopic accumulation of IFT machinery (e.g. kinesin-2 motors, IFT-A, and IFT-B components) in distal cilia segments. Furthermore, retrograde IFT appears to be markedly reduced in the absence of *dyf-5* (Mul et al., 2025). We reasoned that if RIC-8 ciliary trafficking depends on IFT, *dyf-5(ok1177)* mutants may exhibit defective RIC-8 localization. Indeed, RIC-8::TagRFP was detected throughout phasmid cilia, rather than being restricted to the InvC, in 75% of *dyf-5* mutants (Figure 3, C and D). In contrast, only 28% of WT phasmid cilia had any detectable RIC-8::TagRFP in the distal segment (Figure 3, C and D), suggesting that DYF-5 function is important for restricting RIC-8 to the InvC. Notably, *dyf-5* was similarly shown to restrict kinesin-II to the proximal cilium likely by contributing to its undocking from IFT trains (Burghoorn et al., 2007) and to promote unloading of tubulin from IFT complexes at the cilia tip (Jiang et al., 2022). Thus, it would be of interest to test whether DYF-5 also facilitates RIC-8 unloading from IFT trains in the InvC.

### *ric-8* mutants are defective in glycerol responses mediated by ASH neurons

We previously reported that RIC-8 functions as a Gα ODR-3 chaperone in AWC sensory neurons (Campagna et al., 2023). RIC-8 and ODR-3 are also present in the channel cilia of ASH neurons; however, both proteins are largely dispensable for ASH cilia assembly (Roayaie et al., 1998; Salama et al., 2025). Therefore, we next wanted to identify the function of RIC-8 in channel cilia using ASH as a model. First, we tested whether RIC-8 regulates Gα ODR-3 levels in ASH neurons similarly to AWC. To visualize endogenous ODR-3 in ASH cilia, we expressed *wrmScarlet_1-10_* fragment under the control of the *sra-6* promoter in animals that carried *odr-3* endogenously tagged with *wrmScarlet_11_*(ODR-3::split-Sc) (Goudeau et al., 2021; Salama et al., 2025). Consistent with prior work, ODR-3::split-Sc was enriched in WT ASH cilia (Roayaie et al., 1998; Salama et al., 2025) (Figure 4A). Although present inside ASH cilia, ODR-3::split-Sc levels were markedly reduced in strong hypomorphic *ric-8(md1909)* allele compared to WT (Figure 4, A and B), suggesting RIC-8 functions as an ODR-3 chaperone in ASH.

**Figure 4.**
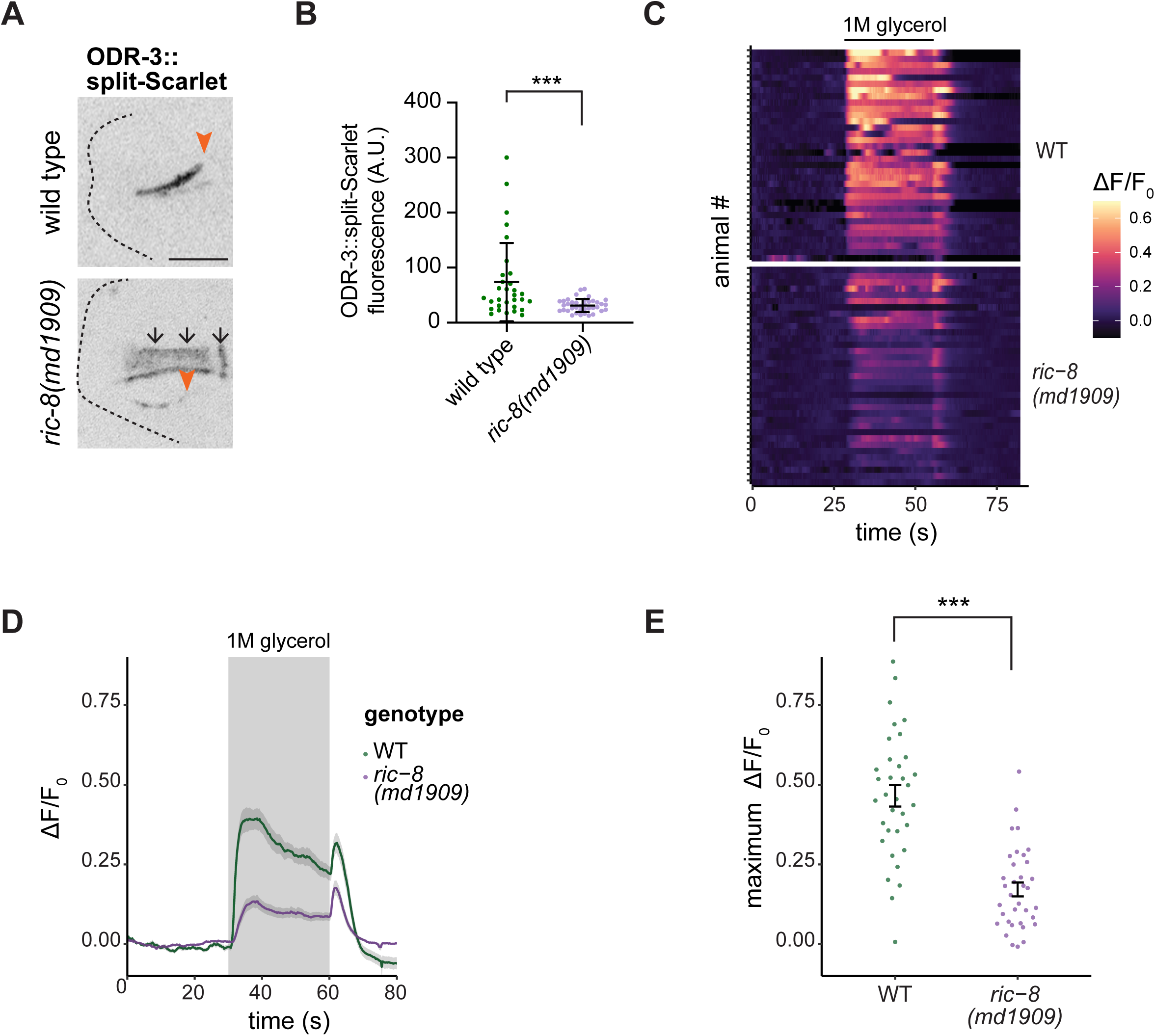
RIC-8 regulates Gα ODR-3 levels and glycerol responses in ASH neurons. (A and B) Images (A) and quantifications (B) of ODR-3::split-Sc in WT and *ric-8(md1909)* ASH neurons. Arrowheads: TZ; arrows: autofluorescence in the pharynx. Scale: 5 μm. *** Different from WT at p<0.001 (Mann-Whitney test). (C) Heatmaps of relative changes in fluorescence intensity (ΔF/F_0_) of GCaMP6 expressed in ASH of the indicated genotypes in response to 1M glycerol. Horizontal bar: glycerol stimulus. Each row in the heatmaps: responses from a single ASH neuron. n=3 days with at least 12 animals/day. (D) Average changes in GCaMP6 fluorescence in ASH for data shown in (C). Shaded regions along the curves: SEM. (E) Quantification of maximum fluorescence intensity change upon glycerol onset in the indicated genotypes. Each dot: the response from a single neuron. *** Different from WT at p<0.001 (Welch’s test).

Gα ODR-3 is a primary transducer of chemosensory signaling in ASH neurons, which mediate avoidance responses to nociceptive chemicals including hyperosmotic solutions such as glycerol (Hilliard et al., 2005; Kato et al., 2014; Yoshida et al., 2012). To test if RIC-8 function is required for cilia-mediated neuronal responses, we examined stimulus-evoked intracellular calcium dynamics in ASH neurons expressing GCaMP6. ASH responses to 1M glycerol were significantly dampened in *ric-8(md1909)* mutants compared to WT (Figure 4, C – E). These responses are similar to those previously reported for *odr-3* mutants (Kato et al., 2014; Yoshida et al., 2012). Thus, our data suggest that RIC-8 functions in ASH to modulate sensory responses likely by controlling ODR-3/Gα levels. Notably, conditional knockout of murine *Ric8b* in olfactory neurons decreased Gα levels and altered olfactory behavior (Machado et al., 2017) akin to our findings in ASH. Although the impact of *Ric8b* deletion on olfactory cilia morphology has not been examined, these findings suggest that RIC-8 function in sensory biology may be evolutionarily conserved.

### Human RIC8A and RIC8B regulate ciliogenesis in RPE-1 cells

Mammalian RIC-8 homologs (RIC8A and RIC8B) have distinct Gα clients. RIC8A functions as a GEF and chaperone toward Gα_i/o_, q, and 12/13, while RIC8B regulates Gα_s/olf_ proteins (Chan et al., 2011; Nagai et al., 2010; Tall et al., 2003; Von Dannecker et al., 2005). To determine if RIC8A and/or RIC8B participate in ciliogenesis, we used small interfering RNAs (siRNAs) to knock down (KD) RIC8A or RIC8B in human RPE-1 cells that ciliate robustly upon serum starvation. Staining cells for ARL13B or acetylated tubulin showed reduced ciliation upon RIC8A and RIC8B KD compared to controls (Figure 5, A and B; Supplemental Figure 2, A and B). We confirmed *RIC8A* and *RIC8B* KD efficiency by qPCR (Figure 5C). Importantly, *RIC8A*-targeting siRNAs did not non-specifically KD *RIC8B* and *vice versa* (Supplemental Figure 2C), and ciliation defects in *RIC8B* KD cells were rescued by co-expression of *RIC8B* cDNA refractory to RNAi using lentivirus (Supplemental Figure 2D). Finally, cells treated with the second set of siRNAs (siRNAs #2) that target *RIC8A* and *RIC8B* coding sequences distinct from those targeted by the first siRNA set (siRNAs #1) resulted in comparable KD efficiency and ciliogenesis defects (Supplemental Figure 2, A, B and E). Collectively, these data indicate that the observed ciliation defects are caused by reduction of RIC8A and RIC8B function.

**Figure 5.**
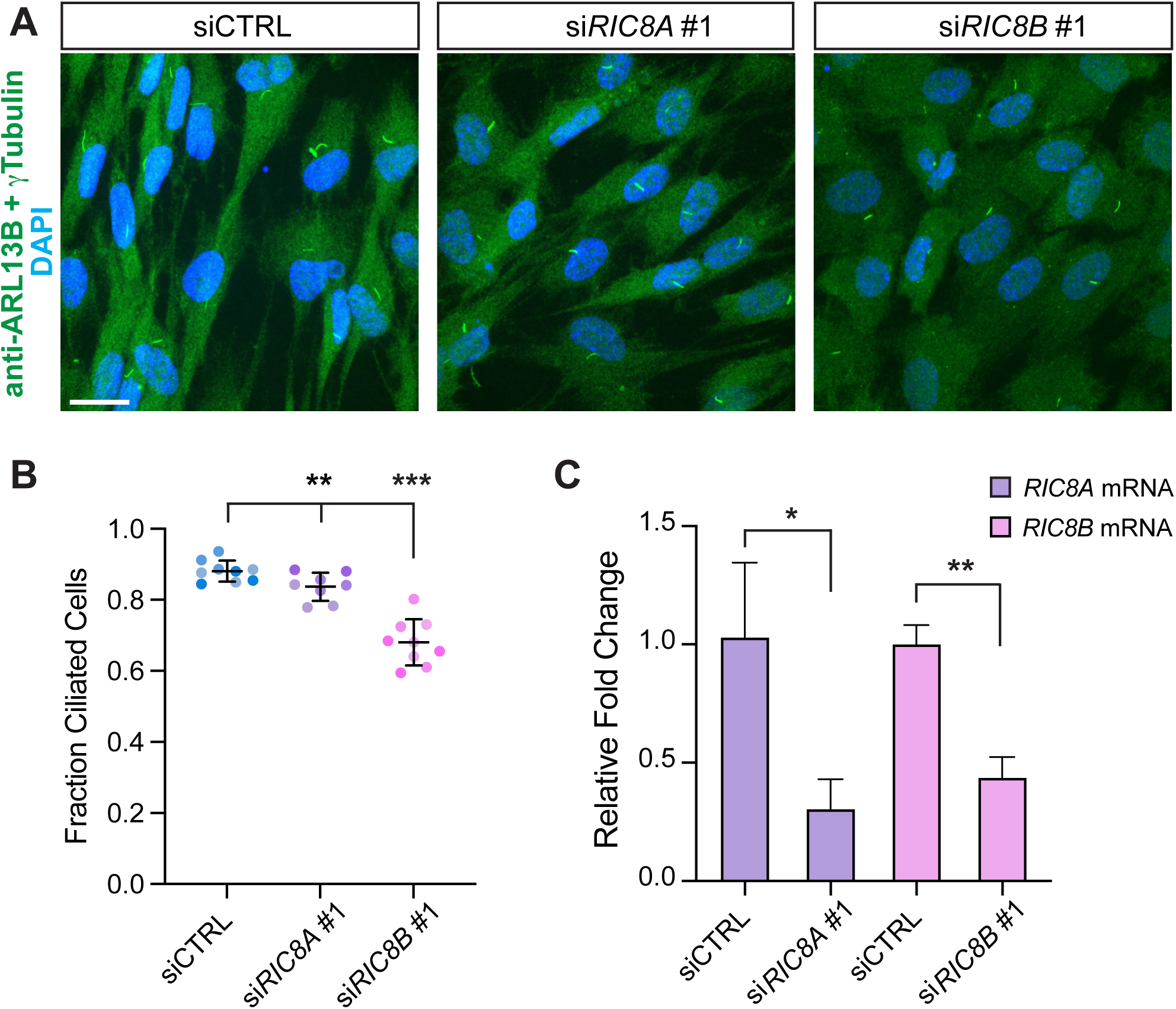
Human RIC8A and RIC8B regulate ciliogenesis in RPE-1 cells. (A) Fixed RPE-1 cells transfected with the indicated siRNAs and stained with the listed antibodies and DAPI. siCTRL: non-targeting siRNA. Scale: 20 μm. (B and C) Quantification of ciliation (B) and relative *RIC8A* and *RIC8B* mRNA levels (C) in RPE-1 cells transfected with the indicated siRNAs. Each data point: one KD experiment; biological replicates are shown in different shades of the corresponding color (B). Summary data in (C) represent 3 biological replicates per condition with 3 technical replicates each. ** and *** Different from siCTRL at p < 0.01 and 0.001, respectively (Fisher’s exact test) (B). * and ** Different between bracketed conditions at p < 0.05 and 0.01, respectively (Welch’s t test) (C).

In summary, our results define trafficking mechanisms that localize *C. elegans* RIC-8 to the InvC, describe a new function for RIC-8 in mediating ASH sensory responses, and demonstrate that human RIC-8 homologs contribute to ciliogenesis in RPE-1 cells, thus highlighting functional versatility of this conserved protein in cilia biology.

## MATERIALS AND METHODS

### C. elegans genetics

*C. elegans* strains were cultured at 20°C on standard nematode growth medium (NGM) seeded with the OP50 strain of *Escherichia coli*. Standard genetic approaches were used to cross in transgenes into mutant backgrounds. All mutant genotypes were confirmed by PCR and/or Sanger sequencing (Azenta). Transgenic *C. elegans* were generated by standard microinjection of DNA and/or ribonucleoprotein complexes into syncytium of hermaphrodite gonad. *The unc-122Δ*p*::gfp* or *unc-122Δ*p*::dsRed* plasmids were used as co-injection markers at 30 and 40 ng/μL, respectively. The same transgenic array was examined in WT and corresponding mutant backgrounds that were directly compared in phenotypic assays.

### CRISPR-Cas9-mediated genome editing

All reagents (crRNA, tracrRNA, single-stranded donor oligonucleotides, and Cas9 protein) were purchased from Integrated DNA Technologies (IDT). CRISPR-Cas9 genome editing to generate the *split-wrmScarlet* (*wrmScarlet_11_*) allele of *ric-8* was carried out as described in (Dokshin et al., 2018). Briefly, the donor oligonucleotide (25 ng/mL), crRNA (56 ng/mL), tracrRNA (100 ng/mL), and Cas9 protein (250 ng/mL) were co-injected with *unc-122D*p*::dsred* co-injection marker into N2 (variety Bristol) WT strain. Transgenic F1 adults were genotyped by PCR and Sanger sequencing (Azenta); F2 individuals homozygous for the transgene were isolated from heterozygous F1 parents to establish transgenic lines. Transgenic strains were outcrossed twice prior to phenotypic analysis.

*ric-8(nch016)* crRNA: 5’ – TCAGAATCCGAATTCTCGGC – 3’

donor oligonucleotide: 5’ – GCCATGTGTTGGAGCTCCTGAAGAATGCTCCAGAACCAGCGCCGGCCGAAAACTCGGATTCTGATGAAGAAGGAGGAGGATCCTACACCGTCGTCGAGCAATACGAGAAGTCCGTCGCCCGTCACTGCACCGGAGGAATGGATGAGTTATACAAGTAATTATTTTTGATTTTTCCATTTTAACATTTTGAAAAAAATTCT – 3’

### Molecular biology

#### Plasmids

The coding sequence corresponding to the RVIP motif of RIC-8 was deleted from the *bbs-8*p::*ric-8^WT^::tagrfp* plasmid (Campagna et al., 2023) by site-directed mutagenesis using QuikChange Lightning kit (Agilent Technologies). The mutagenized construct was verified by full-plasmid sequencing (Plasmidsaurus).

The *RIC8B* rescue plasmid was generated by GenScript. The *RIC8B* cDNA sequence containing five silent mutations in the region targeted by si*RIC8B* siRNA#1 was synthesized and subcloned into a lentiviral vector (*GLV3-CMV-(ORF/RIC8B)-PGK-Puro-P2A-EGFP*) for packaging into lentivirus (GenScript).

#### qPCR

Total RNA was extracted from RPE-1 cells transfected with siControl, si*RIC8A* (siRNA#1 and #2), or si*RIC8B* (siRNA#1 and #2) using the RNeasy kit (Qiagen) per manufacturer’s instructions. RNA samples were reverse-transcribed using the ZymoScript One-Step RT-qPCR Kit (ZymoResearch), and expression levels of *RIC8A* and *RIC8B* were quantified by real-time PCR (Applied Biosystems QuantStudio 6 Pro) relative to *RPL11* control using the 2^-ΔΔCt^ method. Primer sequences are listed below:

*RPL11*: 5’ GTTGGGGAGAGTGGAGACAG 3’ / 5’ TGCCAAAGGATCTGACAGTG 3’

*RIC8A*: 5’ TGATCGCTACTGCTGGAGA 3’ / 5’ TCCAGGGTGAGGAGAACAT 3’

*RIC8B*: 5’ TAGACAGTTGGAAGGTGCATAAA 3’ / 5’ GTCTTCAGTTGGACCTACGATTAG 3’

### Calcium imaging

Young adult worms were transferred to M9 buffer supplemented with poloxamer (1μL/50mL). A single worm was loaded into a microtube using a 3-mL syringe and connected to the olfactory microfluidic chip (Chronis et al., 2007). The inlet channels of the chip were supplied with S-basal buffer and 1M glycerol in S-basal buffer, each connected to computer-controlled rotary valves (Advanced Microfluidics). Recordings were acquired at 10 frames s^-1^, synchronized with the valve switching program. The flow sequence consisted of 30 s of S-basal buffer, followed by 30 s of 1M glycerol, and then 30 s of S-basal buffer for recovery. To correct for photobleaching, an exponential decay was fit to fluorescence intensity values for the first 30 s and the last 20 s of imaging (prior and post stimulus). The resulting curve was subtracted from original intensity values. Amplitude was calculated as maximum change in fluorescence (F-F_0_) in the 10 s following glycerol addition; F_0_ was set to the average ΔF/F_0_ value for 5 s before glycerol onset. Figure panels summarizing calcium imaging data were generated using RStudio.

### RPE-1 cell culture and transfection

Human telomerase-immortalized retinal pigment epithelial cells (hTERT RPE-1) (Nechipurenko et al., 2016) were cultured in DMEM/F12 (1:1) supplemented with 10% fetal bovine serum and 1X antibiotic-antimycotic (Gibco) at 37°C with 5% CO_2_ and tested monthly for mycoplasma using mycoplasma PCR detection kit (ABM). One day prior to transfection, cells were plated in antibiotic-free media at 30,000 cells per well on 12-mm glass pre-treated coverslips (Neuvitro) in a 24-well plate (for immunofluorescence analysis) or at 60,000 cells per well without coverslips in a 12-well plate (for qPCR analysis). Synthetic small interfering RNA oligonucleotides (siRNAs) targeting *RIC8A* or *RIC8B* or non-targeting control siRNA were transfected as previously described (Nechipurenko et al., 2016). For the rescue experiment, RPE-1 cells (untransduced and transduced with *RIC8B*-overexpressing lentivirus) were cultured, plated, and transfected with si*RIC8B* siRNA#1 as described above. The target sequences for siRNAs used in this study are shown below: si*RIC8A* siRNA#1 (J-016121-09-0002, Dharmacon): GGGGAGAUGCUGCGGAACA si*RIC8A* siRNA#2 (J-016121-11-0002, Dharmacon): CAGGAUGCCAUGUGCGAGA si*RIC8B* siRNA#1 (J-021081-09-0002, Dharmacon): UCUCAUCAGUUCCGUGUAA si*RIC8B* siRNA#2 (J-021081-12-0002, Dharmacon): ACAGUUGGAAGGUGCAUAA siControl (D-001810-01-05, Dharmacon): UGGUUUACAUGUCGACUAA

### Lentiviral transduction of RPE-1 cells

RPE-1 cells were seeded at 30,000 cells per well in a 24-well plate and transduced with *RIC8B*-overexpressing lentivirus at a final concentration of 1.8 x 10^6^ units/mL (equivalent to an MOI of 30) for 24 hours in antibiotic-free media containing 8 μg/mL polybrene (GenScript). At 24 hours following transduction, successfully transduced cells were selected with 20 μg/mL puromycin (InvivoGen) for at least 72 hours, or until all untransduced cells in the control well were dead. Expression of *RIC8B* from the lentiviral vector in the transduced cells was confirmed indirectly by the presence of EGFP co-expressed from the same vector as *RIC8B* on an inverted Mateo FL microscope (Leica). Following puromycin selection, *RIC8B*-overexpressing cells were expanded in 60-mm dishes in normal growth media prior to siRNA transfection and immunostaining experiments.

### Immunostaining

RPE-1 cells were fixed in 4% paraformaldehyde for 12 minutes at room temperature (RT) or in ice-cold ethanol or methanol for 10 minutes at -20°C. Fixed cells were blocked in 5% bovine serum albumin (BSA) in phosphate buffered saline with 0.2% Triton X-100 (PBS-T) for one hour at RT or at 4°C overnight and subsequently incubated in primary antibodies diluted in the blocking solution for 1.5 hours at RT or at 4°C overnight. The following primary antibodies were used in this study: anti-ARL13B (1:10, catalog/clone # N295B/66, Developmental Studies Hybridoma Bank), anti-γ-tubulin (1:500, catalog # orb499656, clone # 8D11, biorbyt), anti-acetylated α-tubulin (1:500, catalog # T7451, clone # 6-11B-1, MilliporeSigma). Alpaca anti-mouse Alexa 594 (catalog # 615-584-214, Jackson ImmunoResearch Labs) secondary antibody was diluted in blocking solution and applied for 1.5 hours at RT or at 4°C overnight. DAPI (1:1000, ThermoFisher) was used to stain DNA.

### Microscopy

#### C. elegans

L1 larvae or one-day-old adult hermaphrodites were anesthetized in 10 mM tetramisole hydrochloride (MP Biomedicals) and mounted on 10% agarose pads on top of glass microscope slides. The animals were imaged on an upright THUNDER Imager 3D Tissue (Leica) using 63X NA 1.4-0.60 oil immersion objective and K5 sCMOS camera (Leica) in Leica Application Suite X software. Images of RIC-8::split-Scarlet in Figure 1A and ODR-3::split-Scarlet in Figure 4A were acquired on an inverted Nikon Ti-E microscope with Yokogawa CSU-X1 spinning disk confocal head using 60X NA 1.40 oil immersion objective and ORCA-fusion BT camera (Hamamatsu) in MetaMorph 7 (Molecular Devices). Images for all phenotypic analyses were collected on at least two independent days, and identical acquisition settings were used for imaging all genotypes that were compared directly. In all figures, images are oriented with anterior of the animal to the left.

#### RPE-1 cells

Coverslips with fixed and stained RPE-1 cells were mounted on microscope slides with ProLong Diamond anti-fade mountant (Invitrogen) and imaged on an inverted Nikon Ti-E microscope with a Yokogawa CSU-X1 spinning disk confocal head. Complete z-stacks were acquired at 0.25-μm intervals in MetaMorph 7 software (Molecular Devices) using a 60X NA 1.40 oil immersion objective and an ORCA-Fusion BT Digital CMOS camera (Hamamatsu).

### Image analysis

Image analyses were carried out in Fiji/Image J (National Institute of Health) and are detailed below.

#### RIC-8::TagRFP fluorescence intensity

Fluorescence intensity was quantified by drawing a line from cilia base to the distal tip of the ciliary RIC-8::TagRFP signal and measuring the mean intensity along the line. Similarly, a line was drawn across the PCMC, and the mean intensity along the line was recorded. The relative RIC-8::TagRFP fluorescence for each neuron was reported as the ratio of the mean ciliary intensity over the mean PCMC fluorescence intensity.

#### Line scans

A straight line was drawn from cilia base to the distal boundary of RIC-8::TagRFP signal inside a cilium and measuring fluorescence intensities along the line using the plot profile tool. TagRFP intensity at each point along the line was normalized to the maximum intensity value for that cilium and expressed as percent of the maximum intensity inside the cilium.

#### ODR-3::split-Scarlet fluorescence intensity

The z-slices that encompassed ASH cilia in their entirety were rendered into maximum-intensity projections. Fluorescence intensity of ODR-3::split-Scarlet inside a cilium was quantified by drawing a segmented line from cilium base to tip and measuring the mean intensity along the line after subtracting the average background fluorescence.

### Statistical analyses

Prism 10 software (GraphPad) was used to carry out statistical analyses and generate graphs. In scatter plots, horizontal and vertical bars represent mean ± SD, unless noted otherwise in figure legends. In bar graphs, number of analyzed animals is listed inside corresponding bars. The D’Agostino–Pearson test was used to determine whether the data were normally distributed. Statistical tests and p values are noted in the corresponding figure legends.

## ACKNOWLEDGEMENTS

We are grateful to members of the Nechipurenko lab for critical comments on the manuscript and to Ryan Breitenbach and Thomas L. Morrione for technical assistance and maintenance of strains. Some strains were provided by the CGC, which is funded by NIH Office of Research Infrastructure Programs (P40 OD010440). This work was supported by the NIH (R15 HD109706 – I.N.; R35 GM155316 – I.N.).

