## Supplementary material for "Functions and trafficking mechanisms of RIC-8 in *C. elegans* and mammalian cilia"

**Supplementary Figure 1.** The RVxP motif precludes RIC-8 from localizing to the distal cilium. (A) Alignment of the *C. elegans* RIC-8 (NP 001023561) and human RIC8A (NP 068751) and RIC8B (NP 001317074) sequences showing presence of the RVIP motif (orange box) in the *C. elegans* protein. (B and C) Images (B) and quantification (C) of RIC-8$^{WT}$::TagRFP and RIC-8$^{\Delta RVIP}$::TagRFP localization in phasmid neurons of WT adults. Arrowheads: TZ; arrows: distal boundary of RIC-8::TagRFP signal. d: dendrite. Scale: 5 $\mu$m. * Different from wild type at $p<0.05$ (Mann-Whitney test).

**Supplementary Figure 2.** Ciliation is reduced in *RIC8A* and *RIC8B* KD RPE-1 cells. (A) Immunofluorescence images of fixed RPE-1 cells transfected with the indicated siRNAs and stained with anti-acetylated $\alpha$-tubulin antibody and DAPI. siCTRL: non-targeting siRNA. Scale: 20 $\mu$m. (B – C) Quantification of ciliation (B) and relative RIC8A and RIC8B mRNA levels (C) in RPE-1 cells transfected with the indicated siRNAs. (D) Quantification of ciliation in untransduced RPE-1 cells and RPE-1 cells transduced with *RIC8B*-overexpressing lentivirus after transfection with siControl or si*RIC8B* siRNA #1. Each data point in (B) and (D) represents one KD experiment; biological replicates are shown in different shades of the corresponding color in (D). *** Different from siCTRL (B) or between bracketed conditions (D) at $p<0.001$ (Fisher's exact test). (E) Quantification of relative mRNA levels in RPE-1 cells transfected with the indicated siRNAs. ** Different between bracketed conditions at $p<0.01$ (Welch's t test). Summary data in (C) and (E) represent three biological replicates per condition, with three technical replicates each.

**A**

```
Ce RIC-8   384 I------RVIPPLVSEEVQKRPEENNTLRG 407
Hs RICB8   386 D------QVLPPL--RDVTNRPEVGSTVRN 407
Hs RICBA   354 AQGWPPPQVLPPL--RDVRTRPEVGEMLRN 381
```

**B**

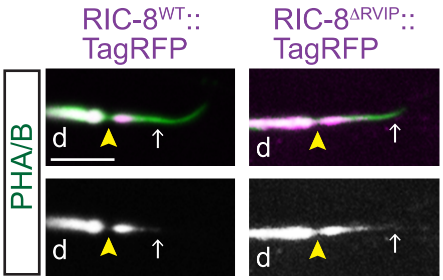


**C**

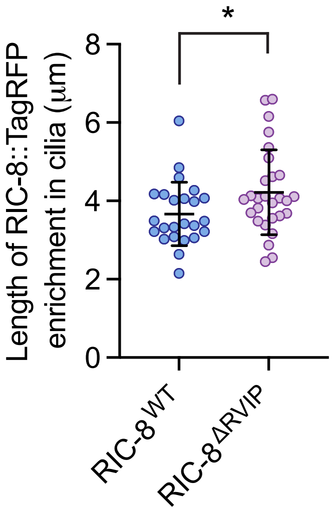

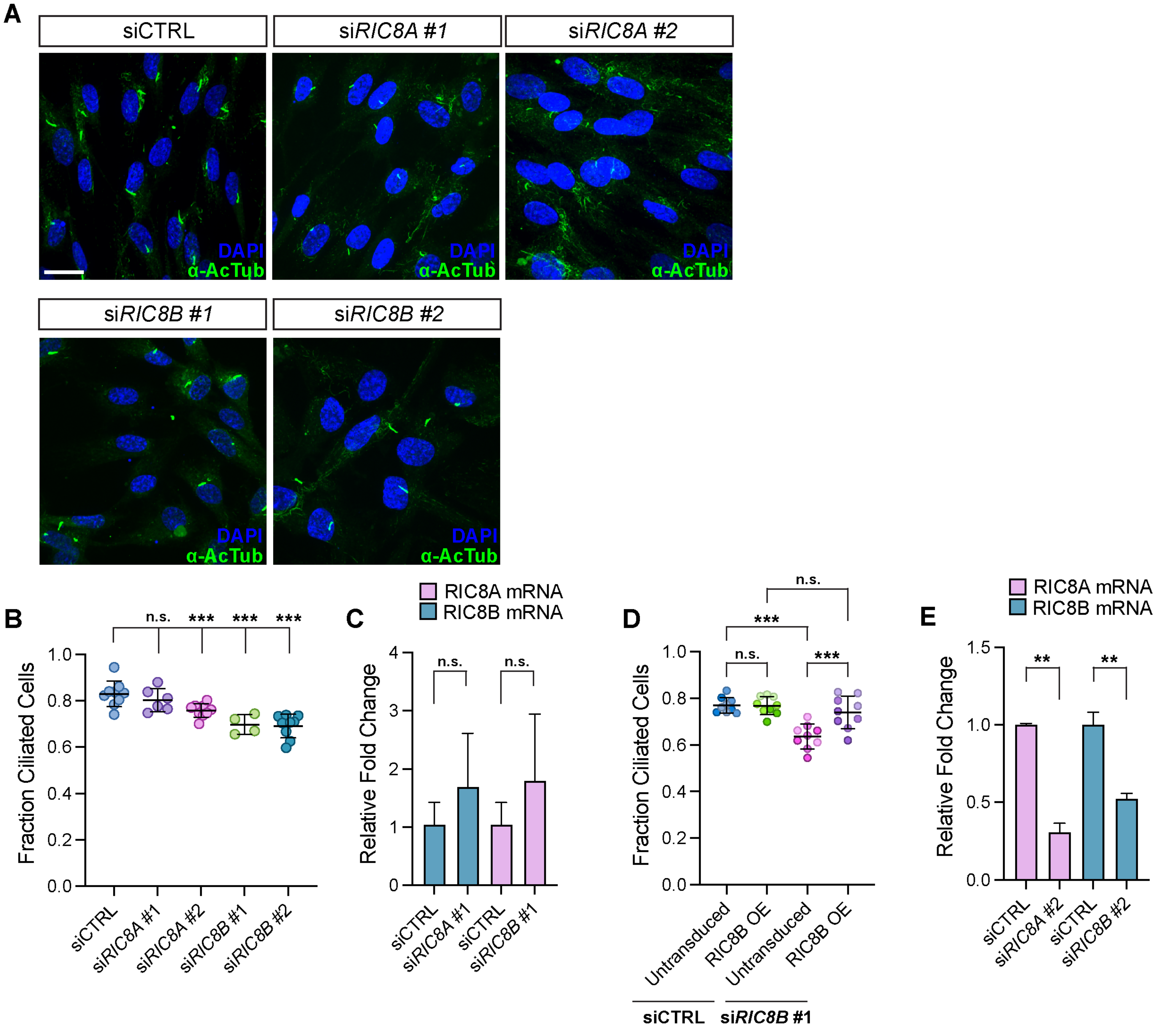
A
siCTRL
siRIC8A #1
siRIC8A #2
siRIC8B #1
siRIC8B #2
DAPI
α-AcTub
B
n.s.
***
Fraction Ciliated Cells
siCTRL
siRIC8A #1
siRIC8A #2
siRIC8B #1
siRIC8B #2
C
RIC8A mRNA
RIC8B mRNA
Relative Fold Change
D
Untransduced
RIC8B OE
siCTRL
siRIC8B #1
E
**

**Supplementary Table 1.** List of strains used in this study

| Strain | Genotype | Source |
|---|---|---|
| NWM740 | ric-8(nch016[ric-8::wrmScarlet$_{11}$]) IV; nchEx018[bbs-8p::wrmscarlet$_{1-10}$, unc-122Δp::gfp] | This work |
| NWM029 | nchEx004[bbs-8p::ric-8::tagrfp, unc-122p::gfp] | This work |
| NWM067 | nphp-2(gk653) V; nchEx004[bbs-8p::ric-8::tagrfp, unc-122Δp::gfp] | This work |
| NWM391 | mks-5(tm3100) II; nchEx002[bbs-8p::myrgfp, bbs-8p::ric-8::tagrfp, unc-122Δp::gfp] | This work |
| NWM331 | kap-1(ok676) II; nchEx002[bbs-8p::myrgfp, bbs-8p::ric-8::tagrfp, unc-122Δπ::gfp] | This work |
| NWM168 | klp-11(tm324) IV; nchEx002[bbs-8p::myrgfp, bbs-8p::ric-8::tagrfp, unc-122Δp::gfp] | This work |
| NWM291 | xbx-1(ok279) V; nchEx002[bbs-8p::myrgfp, bbs-8p::ric-8::tagrfp, unc-122Δp::gfp] | This work |
| NWM310 | che-3(p801) I; nchEx002[bbs-8p::myrgfp, bbs-8p::ric-8::tagrfp, unc-122Δp::gfp] | This work |
| NWM644 | dyf-2(m160) III; nchEx002[bbs-8p::myrgfp, bbs-8p::ric-8::tagrfp, unc-122Δp::gfp] | This work |
| NWM355 | che-11(e1810) V; nchEx002[bbs-8p::myrgfp, bbs-8p::ric-8::tagrfp, unc-122Δp::gfp] | This work |
| NWM639 | daf-10(p821) IV; nchEx002[bbs-8p::myrgfp, bbs-8p::ric-8::tagrfp, unc-122Δp::gfp] | This work |
| NWM616 | ifta-1(nx61) X; nchEx002[bbs-8p::myrgfp, bbs-8p::ric-8::tagrfp, unc-122Δp::gfp] | This work |
| NWM365 | dyf-1(mn335) I; nchEx002[bbs-8p::myrgfp, bbs-8p::ric-8::tagrfp, unc-122Δp::gfp] | This work |
| NWM370 | dyf-13(mn396) II; nchEx002[bbs-8p::myrgfp, bbs-8p::ric-8::tagrfp, unc-122Δp::gfp] | This work |
| NWM628 | dyf-6(m175) X; nchEx002[bbs-8p::myrgfp, bbs-8p::ric-8::tagrfp, unc-122Δp::gfp] | This work |
| NWM319 | ift-74(tm2394) II; nchEx002[bbs-8p::myrgfp, bbs-8p::ric-8::tagrfp, unc-122Δp::gfp] | This work |
| NWM315 | ift-20(ok3191) I; nchEx002[bbs-8p::myrgfp, bbs-8p::ric-8::tagrfp, unc-122Δp::gfp] | This work |
| NWM627 | che-2(e1033) X; nchEx002[bbs-8p::myrgfp, bbs-8p::ric-8::tagrfp, unc-122Δp::gfp] | This work |
| NWM625 | dyf-11(mn392) X; nchEx002[bbs-8p::myrgfp, bbs-8p::ric-8::tagrfp, unc-122Δp::gfp] | This work |
| NWM299 | osm-3(p802) IV; nchEx002[bbs-8p::myrgfp, bbs-8p::ric-8::tagrfp, unc-122Δp::gfp] | This work |
| NWM726 | dyf-5(ok1177) I; nchEx002[bbs-8p::myrgfp, bbs-8p::ric-8::tagrfp, unc-122Δp::gfp] | This work |
| NWM643 | odr-3(nch013[odr-3::wrmscarlet11]) V; nchEx017[sra-6p::wrmscarlet1-10, unc-122Δp::gfp] | This work |
| NWM677 | ric-8(md1909) IV; odr-3(nch013[odr-3::wrmscarlet11]) V; nchEx017[sra-6p::wrmscarlet1-10, unc-122Δp::gfp] | This work |

| | | |
|---|---|---|
| MOS88 | *etyIs1[gpa-13*p*::FLPase, sra-6*p*::FRT::mCherry::Stop::FRT::GCaMP6s]; him-5(e1490) V* | (Pechuk et al., 2022) |
| MOY127 | *etyIs1[gpa-13*p*::FLPase, sra-6*p*::FRT::mCherry::Stop::FRT::GCaMP6s]; ric-8(md1909) IV* | This work |
